# Ionic modulation of interpenetrating network formation, structure and recovery in low-polymeric, supramolecular hyaluronic acid-based hydrogel

**DOI:** 10.1101/2025.09.11.675661

**Authors:** Foluso Akin-Ige, Cindy Rivera, Valentina de Gennaro, Yael Faroud Rivera, Samiul Amin

## Abstract

The formation of interpenetrating polymer networks (IPNs) in polysaccharide-based hydrogels is influenced by ion valency and coordination chemistry. In this work, hyaluronic acid (HA) and kappa-carrageenan (*κ*-CG) were combined at low polymer concentrations suitable for injectable hydrogel applications and enriched with monovalent (K^+^), divalent (Ca^2+^), and trivalent (Al^3+^) ions to investigate ion-specific contributions to network formation, structure and recovery. FTIR spectroscopy showed that K^+^ did not produce detectable sulfate or carboxylate shifts, consistent with a predominantly physical mixture and the absence of IPN formation. Ca^2+^ induced concentration-dependent shifts in both the amide/carboxylate region and the sulfate band (1232 → 1236 cm^−1^), absent at a low concentration but reemerging at higher concentrations, consistent with local chain compaction, complexation of HA carboxylates and bridging between *κ*-CG helices, indicating promotion of a semi-interpenetrated network structure above a threshold concentration. Al^3+^ induced a distinct shoulder in the HA carboxylate region, confirming HA coordination and co-crosslinking with *κ*-CG, yielding a semi-IPN. Microrheology showed progressively stronger local confinement with increasing ion valency corroborating insights from spectroscopy, while recovery tests showed Ca^2+^ attained the highest recovery potential possibly due to more robust network formation, Al^3+^ systems displayed moderate recovery but overscreening at high concentrations resulted in network collapse, and K^+^ systems displayed poor recovery.Importantly, these ion-specific outcomes must be interpreted in the context of combined electrostatic screening effects, which contribute to charge neutralization and polymer chain association independent of coordination chemistry. Collectively, these findings highlight ion charge as critical design levers which may be leveraged in tailoring mechanical properties of interpenetrating network hydrogels for injectable applications.

**Graphical Abstract:** 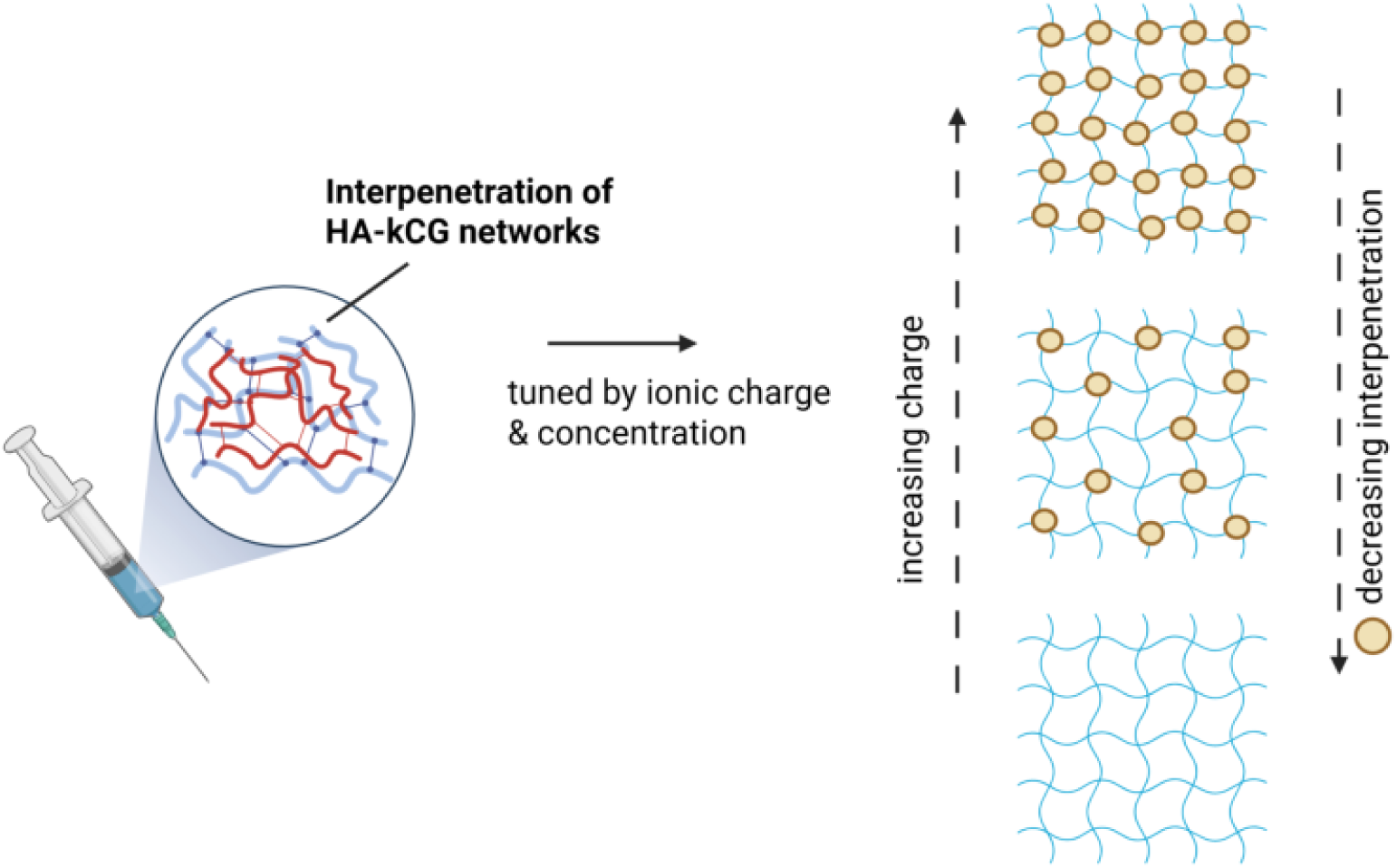

**Highlights:**

- Incorporation of *κ*-carrageenan (*κ*CG) into a chemically unmodified hyaluronic-acid (HA) based hydrogel even at low concentrations is a potentially viable route for more efficient structuring.
- Interpenetrating network formation, structure and recovery of HA– *κ*CG networks can be tuned by charge density and coordination behavior of the crosslinking ion.
- Ca^2+^ reinforces *κ*CG junction zones, yielding the highest elastic moduli and recovery capacity, with evidence of semi-IPN formation pointing to more robust network formation relative to Al^3+^ and K^+^.

## 1. Introduction

Hyaluronic acid (HA), a glycosaminoglycan occurring naturally in the extracellular matrix is widely used in injectable formulations for wound healing applications due to its affinity for cell receptors, high biocompatibility, biodegradability and ability to promote moisture retention, cell attachment, and proliferation [1], [2], [3], [4]. Numerous strategies have been reported for crosslinking HA into hydrogel networks [5], [6], [7],[8]. The most common approach involves covalent crosslinking, which requires chemical modification of the HA backbone with binding sites [9], [10]. Supramolecular HA-based hydrogels synthesized via non-covalent, electrostatic or polyelectrolyte complexation interactions are less commonly explored but are of increasing interest since they are characterized by dynamic, reversible networks that enable stress dissipation and elastic recovery after deformation [11], [12]. Although HA in its native state carries ionizable moieties such as carboxylate, acetamide, and alcohol groups that can, in theory, coordinate with metal ions, earlier studies have shown that divalent cations like calcium or magnesium typically result in only intramolecular charge screening and viscosity reduction rather than gel formation. Cu (II) is the only divalent cation that has been shown to successfully crosslink native HA. Many other studies have often resorted to modifying HA with metal-binding ligands or using multivalent cations such as Fe(III), or Al(III) to achieve effective crosslinking. However, these approaches raise concerns about biocompatibility and/or processing complexity.

A recent study explored the potential of chemically unmodified HA to form supramolecular hydrogels with a wide range of divalent ions beyond Cu^2+^ and identified magnesium (Mg^2+^) as a non-toxic ion capable of creating a new class of Mg-HA hydrogel platforms for biomedical applications [13]. In this study, it was established that effective crosslinking with divalent ions requires high pH conditions to further deprotonate and maximize the negative charges on HA backbone [13]. While there are workarounds for certain systems that could benefit from this approach, the need for high pH introduces additional challenges for wound care applications, particularly when formulating hydrogels to encapsulate sensitive biological molecules like peptides which tend to be notably unstable in highly alkaline environments. Typically, wound healing and controlled release of biologics is enhanced in slightly acidic to neutral pH environments [14].

Carrageenans, a group of naturally occurring, sulfated polysaccharides derived from red algae, are used in many applications to impart structure, stability, stimuli-responsiveness and biocompatibility [15], [16], [17], [18], [19]. Studies have shown that ion valency significantly influences the gelation of carrageenans; kappa- and iota-carrageenan, in particular, are well known for their gelling properties and their ability to form strong yet reversible gel networks in the presence of specific ions [20], [21]. Kappa-carrageenan (*κ*CG) is known to produce stiffer, more brittle gels while iota-carrageenan is known for its softer, more elastic gels but elasticity/stiffness can be tuned in both depending on the application and especially, when combining with other polymers [20]. These dynamic ionic interactions facilitate structural recovery after mechanical deformation. Additionally, in wound healing applications which are a potential use case for our study, carrageenan-based hydrogels have been reported to exhibit anti-coagulant properties, with *κ*CG proving effective in inducing platelet aggregation and fibrin deposition [22], [23], [24]. All of these inform the selection of *κ*CG in the current work.

In this study, we have opted to incorporate *κ*CG into a HA-based hydrogel, hypothesizing that even a simple combination of both polymers will result in a physical mixture with enhanced structure build-up. Some studies have explored this combination in hydrogels although with different design levers [25], [26], [27]. We then leverage the network heterogeneity in this physical mixture by introducing metal ion-polymer interactions, postulating that a possible formation of interpenetrating or semi-interpenetrating networks could occur between both polymers producing independently crosslinked or co-crosslinked networks that exist as physically interlaced networks without covalent linkages. This is a common strategy to enhance mechanical stability of hydrogels. [28], [29], [30], [31], [19], [32], [33]. Additionally, ionic crosslinking is simple, quick, reversible and does not require supplementary initiators and crosslinkers that may induce toxicity in physiological environments [34], [35]. Generally, metal ion-polymer interactions are based on three principles-formation of ion-rich hydrate shell around the biopolymer, formation of complexes via functional groups that allow complexation at neutral pH and formation of complexes at higher alkaline conditions via hydroxyl groups [34]. Our study explores metal-polymer interactions where complexation occurs at neutral pH via sulfate and carboxylic groups on *κ*CG and HA respectively. Since these metal-polymer coordination bonds present a wide range of bonding strengths that can modulate viscoelastic and mechanical properties, we then systematically vary ionic charge and specie, theorizing that as ionic charge increases, the likelihood of HA and *κ*CG forming a semi-interpenetrated network becomes stronger, owing to the ability of higher valency cations to increase electrostatic screening and facilitate multidentate coordination via simultaneous bridging of carboxylate sites on HA and sulfate sites on *κ*CG, as opposed to monovalent ions which bind individually to *κ*CG helices without inducing network formation in HA [36], [26], [37], [38], [39].

Another important consideration of this study is the investigation of structure build-up at low polymer concentrations. Ultra-low to low-polymeric hydrogels are desirable in clinical settings because they are generally easier to inject due to shear-thinning properties and lower high-shear viscosity and can aid clearance of unincorporated material from the body while minimizing risk of inflammatory response [40]. Although there is no universally accepted definition, many studies seem to present an interesting class of hydrogels known as ‘ultra-low polymeric hydrogels’ as systems formed at or below ≤ 1 wt% with some reports demonstrating network formation as low as 0.006 wt%; we identified a study that referred to a 3 wt% hydrogel as a ‘low polymer content system’, though such terminology is not commonly found across literature [41], [42], [43], [44]. Based on this range of usage, we take the liberty of classifying hydrogels prepared with ≈ 1 *<* 3 wt% total solids as low polymer content systems, which is a more fitting description for this study. The combination of all these variables makes our study one that provides novel insight into a potentially viable route to achieve more efficient structuring in and establish design guidelines for chemically unmodified HA-based hydrogel platforms under physiologically relevant pH conditions. To our knowledge, no known study has established this correlation between interpenetrating network formation, structure, and recovery at low polymer concentrations.

## 2. Materials and Methods

### 2.1. Materials

Hyaluronic acid (HA) sodium salt of 400kDa molecular weight was sourced from Making Cosmetics (Washington, USA). Kappa-carrageenan (*κ*CG) was purchased from Sigma-Aldrich (Missouri, USA). Magnesium chloride hexahydrate, sodium chloride, calcium chloride dihydrate, potassium chloride, iron(III) chloride hexahydrate, and aluminum(III) chloride hexahydrate were obtained from Sigma-Aldrich (Missouri, USA). Deionized water with a resistivity of 18 MΩ·cm, obtained from a Thermo Scientific Barnstead E-Pure purification system, was used for all experiments.

### 2.2. HA-κCG Hydrogel Preparation

Sodium hyaluronate and *κ*CG stock solutions were prepared at 5 %w/v and 1.5 %w/v respectively. Stock solutions of sodium chloride, potassium chloride, magnesium chloride hexahydrate, calcium chloride dihydrate, iron(III) chloride hexahydrate, and aluminum(III) chloride hexahydrate were prepared at 1 M concentration and were used as-is without pH adjustments. HA and *κ*CG were then mixed at a volume ratio of 1.25:1 with the total polymer content fixed at 1.8 % w/v in order to remain within low polymer content regime, which has been highlighted as beneficial for injectable hydrogels to maintain ease of administration due to reduced viscosity, faster clearance of unincorporated material and lower risk of inflammatory response. Salt series were prepared at equal molarities (5-75 mM). For each mixture, the appropriate volumes of 1 M salt solution and deionized water required to achieve the desired final ion concentrations at a total volume of 4 mL were added.

### 2.3. Gelation Time Screen

A gelation time screen was conducted at 25°C for all salt series to evaluate time required for gel formation. For each salt, two model species were considered-monovalent (K^+^ and Na^+^); divalent (Mg^2+^ and Ca^2+^) and trivalent (Fe^3+^ and Al^3+^) across concentration range of 5–75 mM. This concentration window was selected because it represents a relevant regime to probe tuning of network assembly and mechanics without exceeding physiologically tolerable osmolality. Similar ranges (tens to hundreds of millimolar) have been widely employed in studies of polysaccharide-based gels/hydrogels to tune gelation kinetics and viscoelasticity [20], [45], [46], [37], [47]. Hydrogels were prepared in glass vials and gelation time was assessed by crude visual observation. The time at which gels did not flow upon inversion was recorded.

### 2.4. Rheology

A rotational rheometer (Netzsch Group, Germany) was used in carrying out extensive rheological characterization. The protocol documented in Chen et al was used, but modified to suit our specific hydrogel system and design considerations, and was performed in triplicate for each sample [48]. Samples were prepared as previously described at 55^◦^C and immediately loaded in sol-form onto a 61 mm stainless steel roughened lower plate geometry and a 40 mm roughened upper parallel plate geometry was lowered to contact the sample at a gap of 1 mm.

A. **Pre-shear** Pre-sheared at shear rate of 0.1 s^−1^ for 10 s to maintain similar sample history across all samples
B. **Temperature ramp** Quenched from loading temperature of 55^◦^C to 25^◦^C at ramp rate of 2^◦^C/min, frequency of 1 Hz and 0.1% strain previously determined from amplitude sweep to know the linear viscoelastic region (LVER) during preliminary rheological characterization of the gels to determine gelation time and temperature.
C. **Time sweep** For 30 mins at 0.1% strain and frequency of 1 Hz to determine plateau moduli.
D. **Heating to 37^°^C** Gels were heated back to 37^◦^C (representing physiological temperature) so subsequent tests could be conducted at this temperature for physiological relevance
E. **Amplitude Sweep with LVER determination** From 0.01 to 100 % strain at 1 Hz frequency, that immediately terminated once instrument detected entry into non-linear viscoelastic region to preserve structure of gel for subsequent tests that rely on accurate moduli values.
F. **Time sweep** Conditioning time of 10 mins at 0.1% strain and frequency of 1 Hz to stabilize gel and determine plateau moduli at 37^◦^C.
G. **Frequency Sweep** From 0.1 to 10 Hz at 0.1% strain.
H. **High Strain Sweep** For 1 min at 100 strain and frequency of 1 Hz
I. **Low Strain Sweep** For 15 mins at 0.1 strain and frequency of 1 Hz to determine recovery rate
J. **Cyclic Sweep Time sweeps** of 1 min at 100% strain, 1 Hz and 2 min at 0.1% strain, 1 Hz in 3 cycles.

### 2.5. Diffusing Wave Spectroscopy (DWS)

DWS experiments were carried out with a diffusing wave spectrophotometer (LS Instruments AG, Switzerland) based on echo-DWS technique for non-ergodic media and were performed twice for each sample. Gel samples were prepared at 55^◦^C using the method described previously and 1 %v/v of 600 nm polystyrene tracer particles obtained as a 10 wt% suspension (Sigma Aldrich, Missouri) were added to the gel samples. 2 mL of the sample was then immediately pipetted into a 10mm path length glass cuvette and loaded into the instrument.

### 2.6. Fourier-transform Infrared (FTIR) Spectroscopy

4 mL gel samples were prepared as previously described and transferred into glass petri dishes to air-dry for 48 hours, after which the dried thin films obtained (xerogels) were subsequently studied using the ATR–FTIR technique. All reported FTIR spectra were collected using a Perkin Elmer Spectrum-100 FTIR spectrophotometer at room temperature within the range of 600–4000 cm^−1^ with 16 scans.

### 2.7. Data analysis

Data cleaning and preprocessing were performed in Microsoft Excel, statistical analyses were conducted in R, and all other analyses were carried out using OriginPro 2025b.

## 3. Results and Discussion

### 3.1. Gelation time screen

The hydrogels shown in Fig. 1 appear similar, but Fig. 2 shows a simple heatmap classifying their ion- and concentration-specific behavior into ‘slow’, ‘moderate’, and ‘fast’ gelation. To minimize bias, this classification was derived objectively using a combination of hierarchical clustering and principal component analysis (PCA) shown in Figs. S1 and S2 to identify underlying patterns in the gelation time screening data. Both PCA and hierarchical clustering consistently classified Na^+^ as the slowest gelator and Mg^2+^ as intermediate, while K^+^, Ca^2+^, Fe^3+^, and Al^3+^ formed the ‘fast’ gelator class. PCA emphasized the stronger separation of Fe^3+^ and Al^3+^ from the rest along PC1, highlighting their rapid gelation profile, whereas hierarchical clustering revealed low cophenetic distances to K^+^ and Ca^2+^, suggesting that all four ions share broadly similar gelation profiles. These methodological differences account for minor discrepancies in the intermediate ordering, but the consensus classification across both analyses yields the rank order:

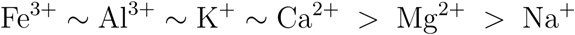

**Figure 1:**
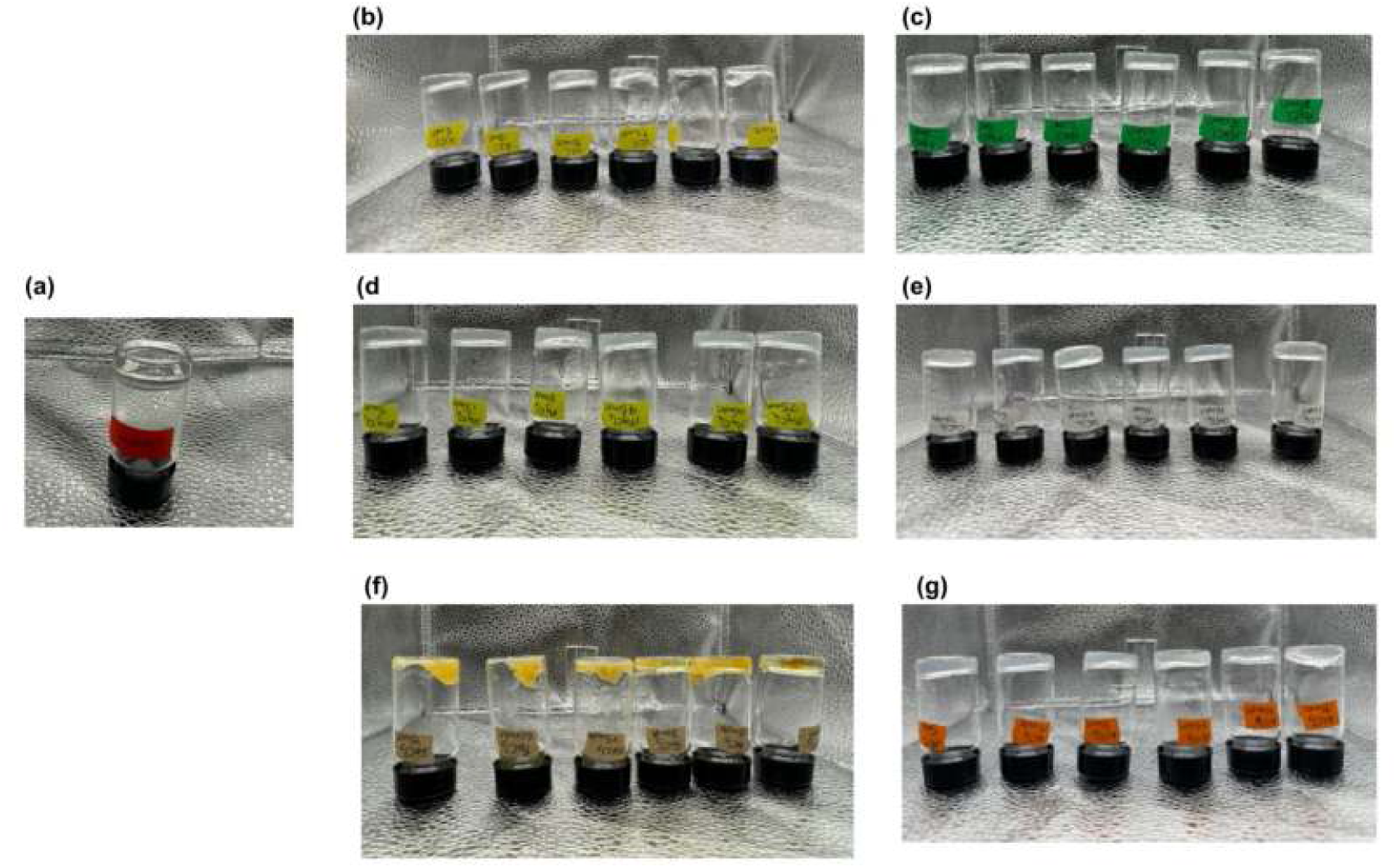
Inverted vials showing blank HA-kCG gel (a), and HA-kCG gels in which 5-75 mM of KCl (b), NaCl (c), MgCl_2_ (d), CaCl_2_ (e), FeCl_3_ (f), and AlCl_3_ (g) were added

**Figure 2:**
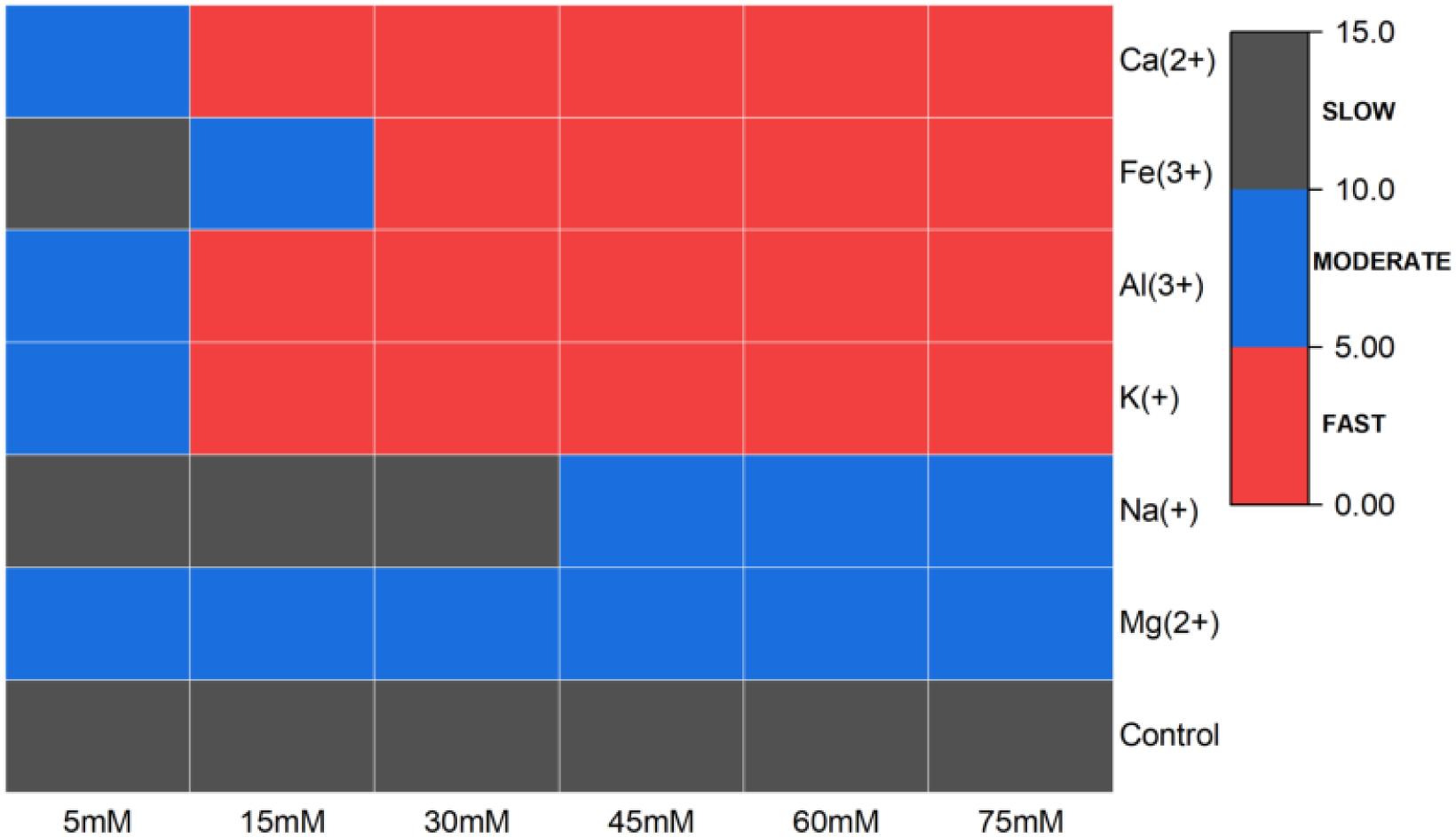
Heatmap showing gelation time screen

In the case of the monovalent ions, both Na^+^ and K^+^ promote *κ*CG macroscopic gelation due to decreased electrostatic repulsions which result in sulfate charge screening and increased association between the chains [15]. The propensity of K^+^ to induce a coil to double helix transition preceding and promoting aggregation in *κ*CG while yielding stronger, more ordered gels has been well documented [39]. Na^+^, on the other hand, even though it does not promote the coil to double helix transition, takes place in the aggregation process and forms weak gels with more disordered structure relative to that obtained with K^+^[39]. We see that Na^+^ displays significant concentration-dependent behavior as well, gelling the mixture faster above 30mM aligning with trends from other studies that *κ*CG responds to Na^+^ at higher concentrations. [39]. In contrast to *κ*CG, Sheehan et al’s study elucidated on the effect of monovalent ions on hyaluronic acid in solutions around pH 6-they only yield pairwise ‘binding on touching’ of hyaluronate segments, they do not promote significant interpenetration of hyaluronate chains hence, they are not capable of inducing continuous network formation in HA polymer [37]. In a study that was conducted at higher pH conditions, a range of monovalent ions beyond K^+^ and Na^+^ were not able to interact with the carboxylate groups even in a strongly deprotonated state, regardless of ionic radius [13]. It is thus clear that in the hydrogels formed with monovalents, HA cannot be a partaker in network formation; *κ*CG drives gelation and since it responds to K^+^ more readily, the network is formed fastest, relative to Na^+^.

With the multivalent ions, it is pertinent to note that network formation must be interpreted in light of both increased electrostatic screening and potential direct coordination to HA and *κ*CG functional groups [45], [49]. This study was designed around equimolar concentrations of the salt series to reflect practical dosing scenarios however, ionic strength does not scale linearly with molarity when ions of different valencies are compared. This scaling arises from the general expression for ionic strength,

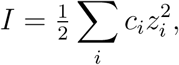

which is derived from the Debye–Hückel theory of electrolytes, specifically from the expression for the inverse Debye length,

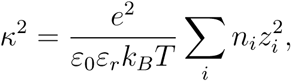

where *c_i_* (or *n_i_*) is the ion concentration, *z_i_* is the ionic valency, and the *z*^2^ term amplifies the contribution of multivalent ions to screening [45], [49]. This then means that at the same molar concentration, divalent salts contribute roughly threefold ionic strength, while trivalent salts deliver about sixfold higher ionic strength than monovalent salts. Nevertheless, coordination is still likely. Among the divalent ions, Ca^2+^ is able to yield gels faster than Mg^2+^. In theory, Ca^2+^ is able to truly bind and facilitate interpenetration of hyaluronate chains. This may lead us to believe that unlike monovalents, HA forms a network here but it has been established that even though ion-HA complexes are formed in the presence of divalents like Ca^2+^ and Mg^2+^, this does not lead to formation of a space-spanning network [38]. In fact, at higher pH, Mg^2+^ is able to crosslink unmodified HA where Ca^2+^ is not, likely due to its smaller ionic radius relative to Ca^2+^, where its larger ionic radii can affect ion-HA bond stability or pose steric hindrances to crosslinking [13]. Thus, we turn our attention to *κ*CG. Divalent cations like Ca^2+^ and Mg^2+^, can indeed facilitate coil-to-helix transition and aggregation of *κ*CG helices since their suggested binding mechanism entails binding directly between carrageenan helices and creating more junction zones in resulting gels rather than binding to single sulfate groups on *κ*CG helices as with monovalents [45]; the order of influence also suggests that Ca^2+^ is able to link the carrageenan helices together more readily than Mg^2+^ [50], [51]. Ca^2+^-sulfate contact pairing is thermodynamically more favorable hence, direct coordination to sulfate sites on CG can occur, producing bi-/multidentate geometries that bridge helices more efficiently. Mg^2+^, by contrast, exhibits a slower response to bridging because it prefers to remain in solvent-shared pairs 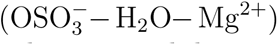 [52]. Thus, with divalents, it is more likely that *κ*CG is the principal driver in network formation with Ca^2+^ exerting a stronger influence, albeit via a different mechanism that may allow for trapping of HA polymer chains within the *κ*CG network.

With trivalents, both Fe^3+^ and Al^3+^ form gels rapidly, but produce different network architectures. Fe^3+^ was observed to form inhomogenous gels (Fig. 1f) exhibiting syneresis unlike Al^3+^ in which bulk gel formation was uniform at all concentrations. Fe^3+^ is regarded as a ‘hard’ metal cation that tends to form complexes leading to formation of 3-D networks in polysaccharide ligands such as alginates, carrageenans, HA and even chitosan [53], [54], [55], [56]. With HA, complexation occurs via coordination with negatively charged carboxyl groups yielding mono-, bi- and tri-dentate bonds depending on pH and ligand availability [54], [57]. Its interesting binding character often confers desirable mechanical properties in many instances however in many bulk hydrogel systems that crosslink with Fe^3+^, free Fe^3+^ is hardly used and often Fe-EDTA complex is used to control the physical crosslinking process to give homogenous bulk gels for injectable applications [26], [58], [59], [60]. An invention disclosed the formation of a homogenous gel with pH close to neutral by using Al^3+^ to crosslink hyaluronic acid. Herein, it was noted that due to strong ionic bonds created with ferric chloride, it is necessary to adjust pH of HA to acidic pH before mixing and so, there is the added processing step to achieve homogenous gel formation with ferric chloride unlike with aluminium chloride [61]. Al^3+^ also interacts strongly with *κ*CG’s sulfate esters and can act as a multidentate node bridging *κ*CG helices, increasing junction zone density and resulting in rapid gelation. A study developed high-tensile-strength carrageenan fibers in which coordination and ionic bonds between Al^3+^, -OH and sulfate groups facilitated crosslinking [62]. With the trivalents, a co-crosslinked network between HA and *κ*CG is more likely since both polymers can participate in network formation via their carboxylate and sulfate sites respectively.

With the above insights gained from the gelation screen on ion- and concentration-specific trends, K^+^, Ca^2+^ and Al^3+^ were selected for further studies at equimolar concentrations of 15, 30 and 60 mM on interpenetrating network formation, structure and recovery in HA-*κ*CG hydrogel, with Al^3+^ being the representative ion in trivalent category to ensure that all ions were studied under the same conditions without introducing additional processing steps different from those used for the other gels.

### 3.2. Interpenetrating Network Formation

In this section, we address the possibility of HA and *κ*CG forming an interpenetrating network hydrogel, the suggested type of interpenetrating networks formed and how ionic charge and concentration modulate the formation of this interpenetrating network, combining insights from FTIR spectroscopy, rheology and microrheology.

#### 3.2.1. FTIR Spectroscopy

In Fig. 3a, *κ*CG showed characteristic sulfate and anhydrogalactose bands at 1222, 927, and 840 cm^−1^, while HA exhibited distinct amide I/II vibrations around 1603 cm^−1^ and a carboxylic acid C–O–H bending mode around 1411 cm^−1^ [26], [25]. When the physical mixture of HA and *κ*CG were compared to their ion-enriched counterparts in Figs. 3b to 3d, spectral shifts were observed with differing intensities via ionic charge (Table 1).

**Figure 3:**
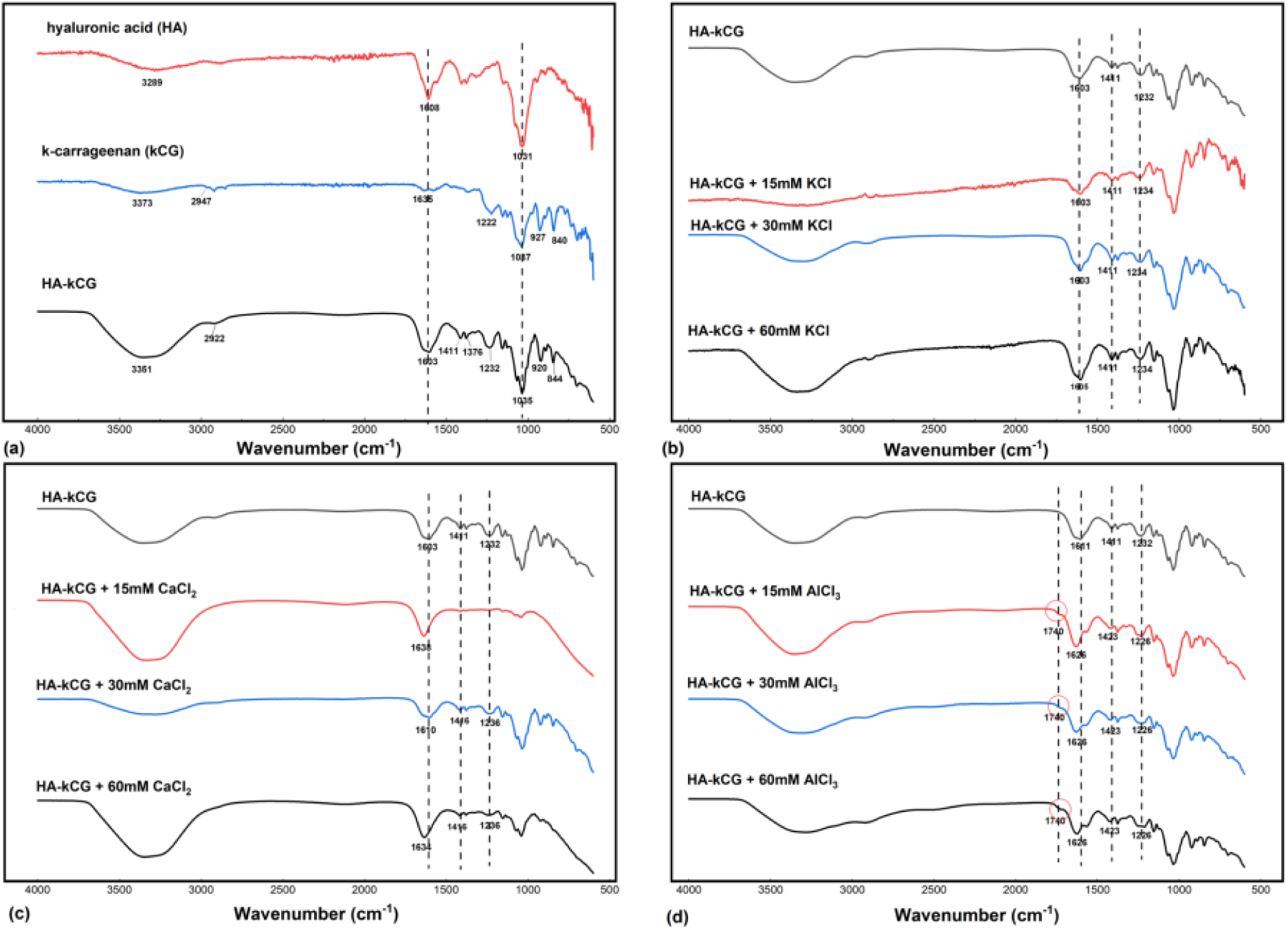
FTIR spectra of pure HA and `κ`CG and HA-`κ`CG physical mixture (a), HA-`κ`CG with 15, 30 and 60 mM KCl (b), HA-`κ`CG with 15, 30 and 60 mM CaCl_2_ (c), and HA-`κ`CG with 15, 30 and 60 mM AlCl_3_

**Table 1:** Spectral shifts in HA–κCG system.

| Sample | Amide region<br>(1600–1620 $\text{cm}^{-1}$) | SO<sub>4</sub> asymmetric<br>stretching<br>(1100–1250 $\text{cm}^{-1}$) | C–O–H bending of<br>carboxylate<br>($\sim 1450 \text{ cm}^{-1}$) | Carboxylate<br>coordination peak<br>($\sim 1700 \text{ cm}^{-1}$) |
| --- | --- | --- | --- | --- |
| HA-κCG | 1603 | 1232 | 1411 | – |
| HA-κCG + 15 mM KCl | 1603 | 1234 | 1411 | – |
| HA-κCG + 30 mM KCl | 1603 | 1234 | 1411 | – |
| HA-κCG + 60 mM KCl | 1603 | 1234 | 1411 | – |
| HA-κCG + 15 mM CaCl<sub>2</sub> | 1638 | – | – | – |
| HA-κCG + 30 mM CaCl<sub>2</sub> | 1610 | 1236 | 1416 | – |
| HA-κCG + 60 mM CaCl<sub>2</sub> | 1634 | 1236 | 1416 | – |
| HA-κCG + 15 mM AlCl<sub>3</sub> | 1623 | 1226 | 1423 | 1740 |
| HA-κCG + 30 mM AlCl<sub>3</sub> | 1626 | 1226 | 1423 | 1740 |
| HA-κCG + 60 mM AlCl<sub>3</sub> | 1626 | 1236 | 1423 | 1740 |

HA and *κ*CG are known to interact primarily through hydrogen bonding [26], and with increasing KCl concentration, changes in the intensity and broadness of the -OH band were observed, suggesting interactions via hydrogen bonding. In the sulfate ester region, the band at 1232 cm^−1^ in HA–*κ*CG alone shifted slightly to 1234 cm^−1^ upon KCl addition at all concentrations. Such a minor shift does not constitute a true chemical shift and seems to agree with previous work on *κ*CG ‘crosslinked’ with KCl, where no significant spectral changes were reported with KCl-enriched carrageenan beads [63]. This suggests that monovalent cations do not induce crosslinking or interpenetrating network formation but rather support a predominantly physical mixture. This interpretation is further supported by the absence of detectable spectral shifts in the amide/carboxylate region (around 1600 cm^−1^ and around 1450 cm^−1^), indicating no binding of K^+^ to HA carboxylate groups [37]. By contrast, the CaCl_2_ series exhibited shifts in the amide/carboxylate region and a more pronounced sulfate shift (1232 to 1236 cm^−1^), demonstrating that even though no gel formation occurs with CaCl_2_ at neutral pH, spectroscopy is able to capture the structural signatures that seem attributable to local chain compaction and binding of HA carboxylates, while also capturing the distinct mechanism of Ca^2+^ bridging between *κ*CG helices rather than the mechanism of K^+^ exhibiting individual ion pairing at sulfate sites which may explain why no significant shift in sulfate stretching band was detected with KCl[45],[37]. Notably, at 15mM CaCl_2_, both sulfate and carboxylate bands were undetectable, likely due to low ionic concentration. As concentration increased, both regions re-appeared with measurable shifts supporting increased ordering. With these spectroscopic insights, our proposed interpretation is that the likelihood of HA chains to become entrapped within *κ*CG junction zones is increased, promoting a semi-interpenetrated structure but only above a certain concentration threshold. With AlCl_3_, a distinct shoulder appeared in the HA carboxylate region near 1740 cm^−1^ at all concentrations. This feature, absent in the physical mixture and in the K^+^- or Ca^2+^-enriched gels, points to COO^−^ coordination capable of promoting HA crosslinking. This interpretation is consistent with the findings of Mashaqbeh et al. who reported a comparable shoulder at 1735 cm^−1^ arising from Fe^3+^ coordination to HA [26]. Given the chemical similarity of Al^3+^ and Fe^3+^, a similar coordination mechanism is plausible here. Additional support for Al^3+^ binding comes from shifts in both the amide and carboxylate bands, as well as in the sulfate stretching region, suggesting that Al^3+^ can simultaneously bridge HA carboxylates and *κ*CG sulfates. Taken together, these results support the formation of a co-crosslinked semi-interpenetrating network (semi-IPN), in which Al^3+^ ions bridge junction zones across both polymers. However, because the system is prepared in a one-pot synthesis, it is unlikely to yield two independently crosslinked subnetworks, and thus does not represent a fully interpenetrated network [64]. Thus, at best, spectroscopic insights point to promotion of semi-interpenetrated network structure as ion charge increases.

#### 3.2.2. Microrheology

MSD data shown in Fig. 4 further supported these findings, showing reduced probe mobility with increasing ion valency. Data were obtained and shown for 30 mM systems, where the order of confinement followed: no ion *>* K^+^ *>* Ca^2+^ ≈ Al^3+^. The lowest MSD values were observed for Al^3+^, nearly overlapping with Ca^2+^, consistent with stronger local confinement and suggesting that semi-interpenetrated structures restrict probe mobility more effectively than the physical mixture formed in the case of K^+^. In the K^+^ system, the MSD curve was slightly higher, reflecting insufficient junction formation which will allow probes to diffuse more freely.

**Figure 4:**
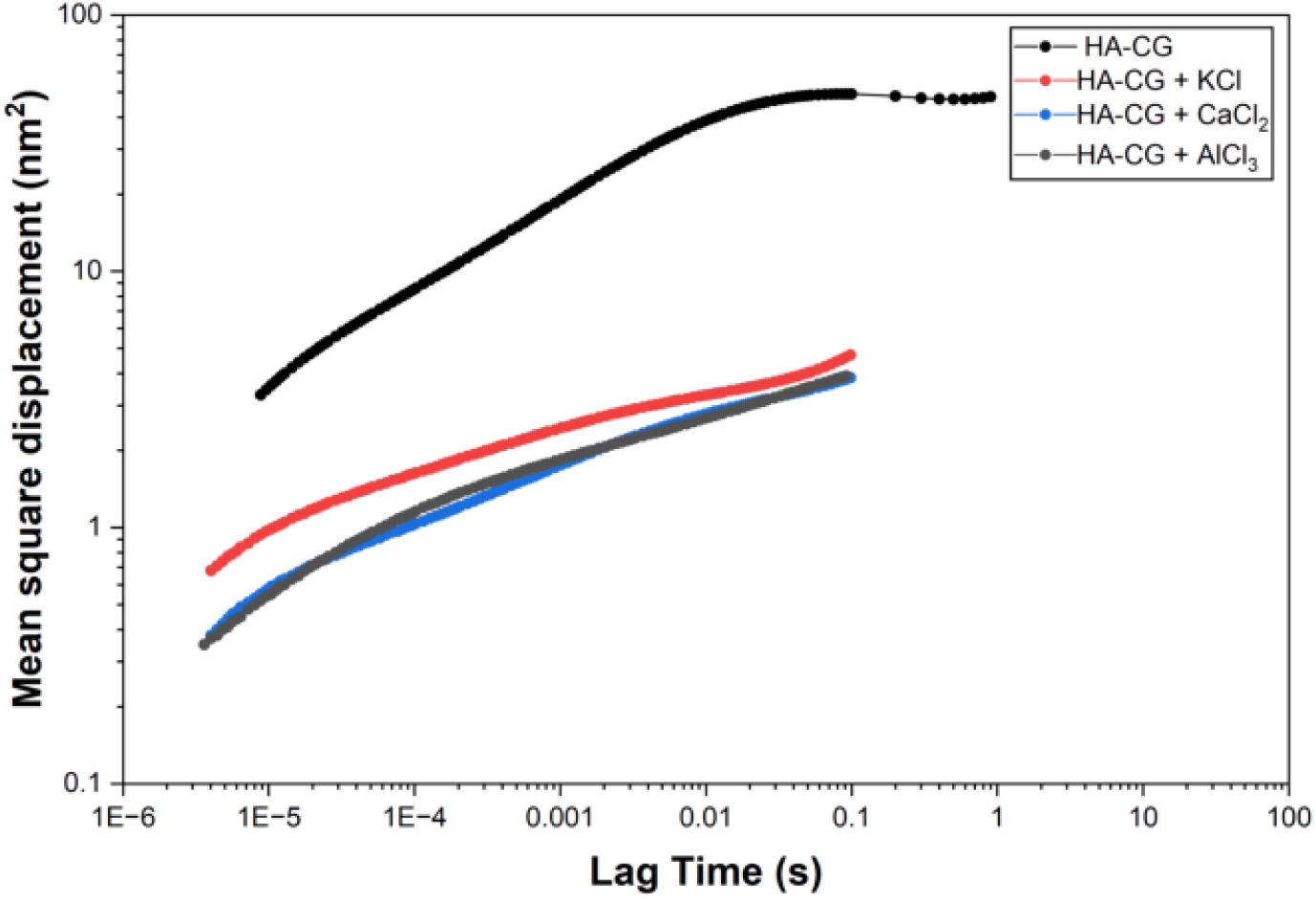
Mean square displacement by ion charge

#### 3.2.3. Bulk rheology

We theorized that combining HA with *κ*CG would enhance network structure beyond what either polymer achieves individually. This was confirmed by comparing elastic moduli obtained via small-amplitude oscillatory shear characterization at 0.1% strain, where the storage modulus (*G*^′^) of the HA–*κ*CG blend increased by nearly two orders of magnitude compared to HA alone and by more than one order of magnitude compared to *κ*CG alone, demonstrating strong synergistic effect on structure build-up (Fig. 5a). Additionally, a narrowing of the linear viscoelastic region when *κ*CG was incorporated although not as pronounced as that of the ion–HA–*κ*CG systems, confirmed this theory of synergistic structural enhancement with incorporation of *κ*CG and subsquently, in the presence of ions (Table 2). Frequency sweeps (Fig. S3) further established that all systems displayed solid-like behavior (i.e., *G*^′^ ≫ *G*^′′^), with minimal frequency dependence of *G*^′^ in nearly all systems. In the representative frequency sweep plot shown in Fig. 5b, we see that the loss tangent (tan *δ* = *G*^′′^*/G*^′^) followed the order: Ca^2+^ *>* K^+^ ≈ Al^3+^-enriched, possibly indicating the more elastic nature of Ca^2+^-enriched networks compared to other ions. To obtain elastic moduli plateau 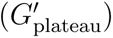 under the different conditions considered in this study, the plateau region in the time-sweep curves presented in Figs. S4, S5 and S6 were generally identified as the region that satisfied the condition in Equation 1 (typically the last 300 s of the time-sweep) and G’ values in this region were then averaged to obtain a single plateau value;

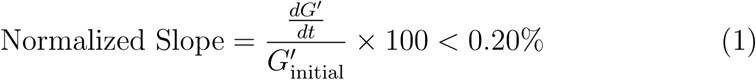

**Figure 5:**
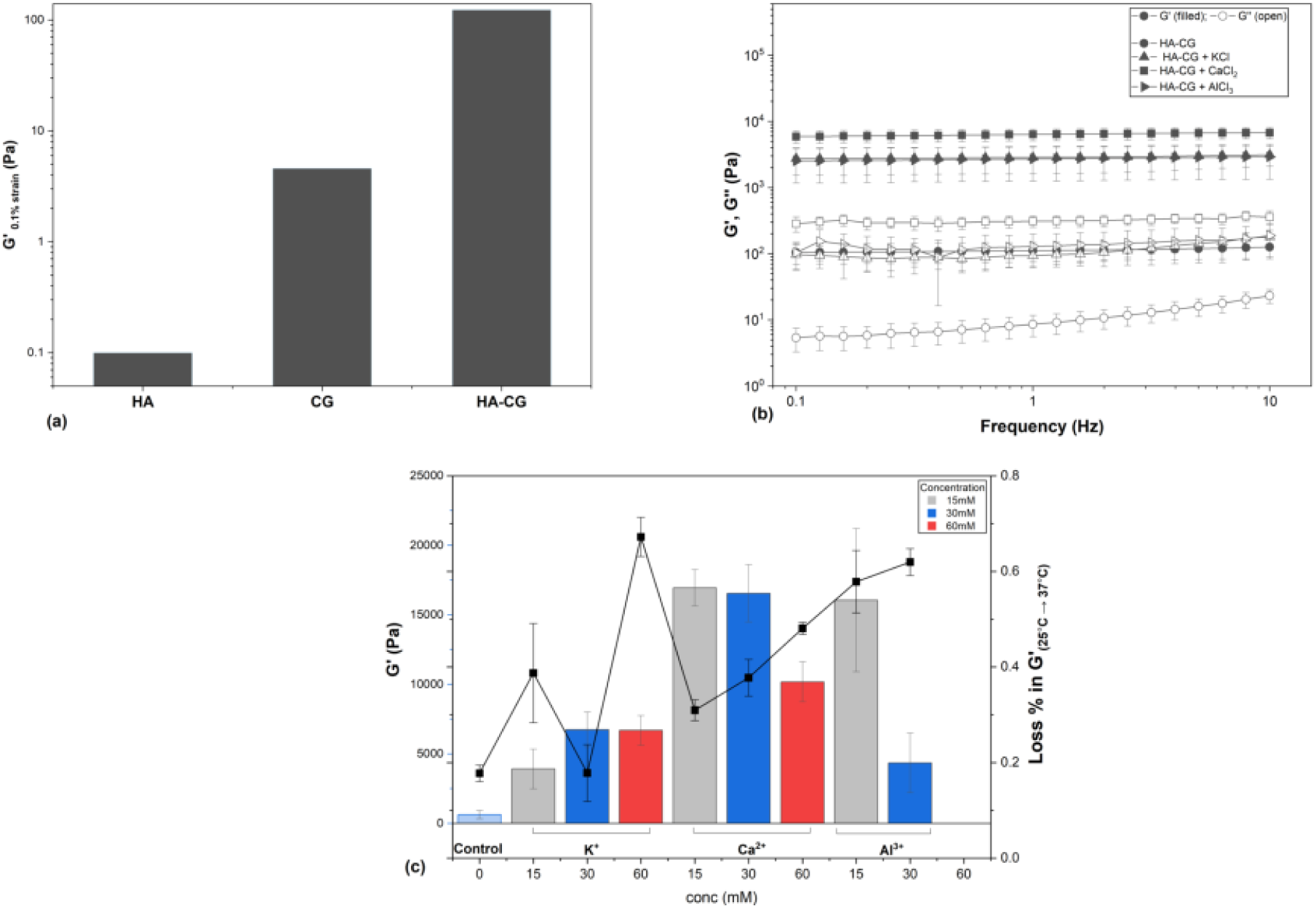
Rheology plots showing elastic moduli at 0.1% strain for HA, `κ`CG and HA-`κ`CG (a), frequency sweeps for representative HA-`κ`CG gels at 30 mM ion concentration (b), and elastic moduli at 25C shown with loss % in elastic moduli on heating gels from 25^◦^C to 37^◦^C (c)

**Table 2:** Linear Viscoelastic Region (LVR) ranges for different HA–κCG and ion-containing systems.

| Samples | LVR (%) |
| --- | --- |
| HA | 0.1–18.64 |
| CG | 0.1–13.90 |
| HA-κCG | 0.1–15.86 |
| HA-κCG + 15 mM KCl | 0.01–3.29 |
| HA-κCG + 15 mM CaCl<sub>2</sub> | 0.01–3.26 |
| HA-κCG + 15 mM AlCl<sub>3</sub> | 0.01–0.91 |
| HA-κCG + 30 mM KCl | 0.01–7.82 |
| HA-κCG + 30 mM CaCl<sub>2</sub> | 0.01–2.34 |
| HA-κCG + 30 mM AlCl<sub>3</sub> | 0.01–1.33 |
| HA-κCG + 60 mM KCl | 0.01–0.52 |
| HA-κCG + 60 mM CaCl<sub>2</sub> | 0.01–1.96 |

From Fig. S4 and Fig. S5, elastic moduli plateau at 37^◦^C was compared across all ion conditions and elastic moduli at 37^◦^C was computed as loss percent of elastic moduli plateau at 25 °C (Fig. 5c) using Equation 2;

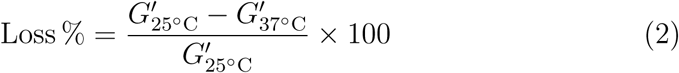

This was in a bid to understand the structural behavior of the gels after having been injected at physiological temperature, the trend observed was again, similar to that of the loss tangent where Ca^2+^-enriched network displayed relatively lower loss % in elastic moduli compared to the other ions.

### 3.3. Network Recovery

Based on the insights in previous section, we will now address how the formation of IPN influences network recovery in our HA–*κ*CG system. Recovery after deformation provides insight into how well the HA-kCG networks can recover their structure, which may be correlated to whether the systems behave as independent networks or show features of semi-IPN formation. The G’ plateau from the plots presented in Fig. S6 were expressed as a percentage of initial G’ (at a working temperature of 37^◦^C) to determine recovery % using Equation

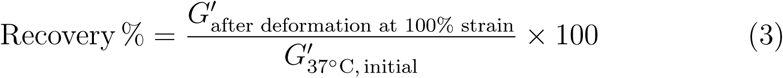

We already see that Ca^2+^ networks point to a more robust formation, this was further reflected in the recovery profile presented in Fig. 6. Statistical analyses (Section 2 of Supporting Information) revealed the recovery values depended strongly on both ion valency and concentration (MANOVA: *F* = 11.09*, p <* 0.001 for valency; *F* = 9.08*, p <* 0.001 for concentration), with a significant interaction effect (*F* = 7.44*, p <* 0.001).

**Figure 6:**
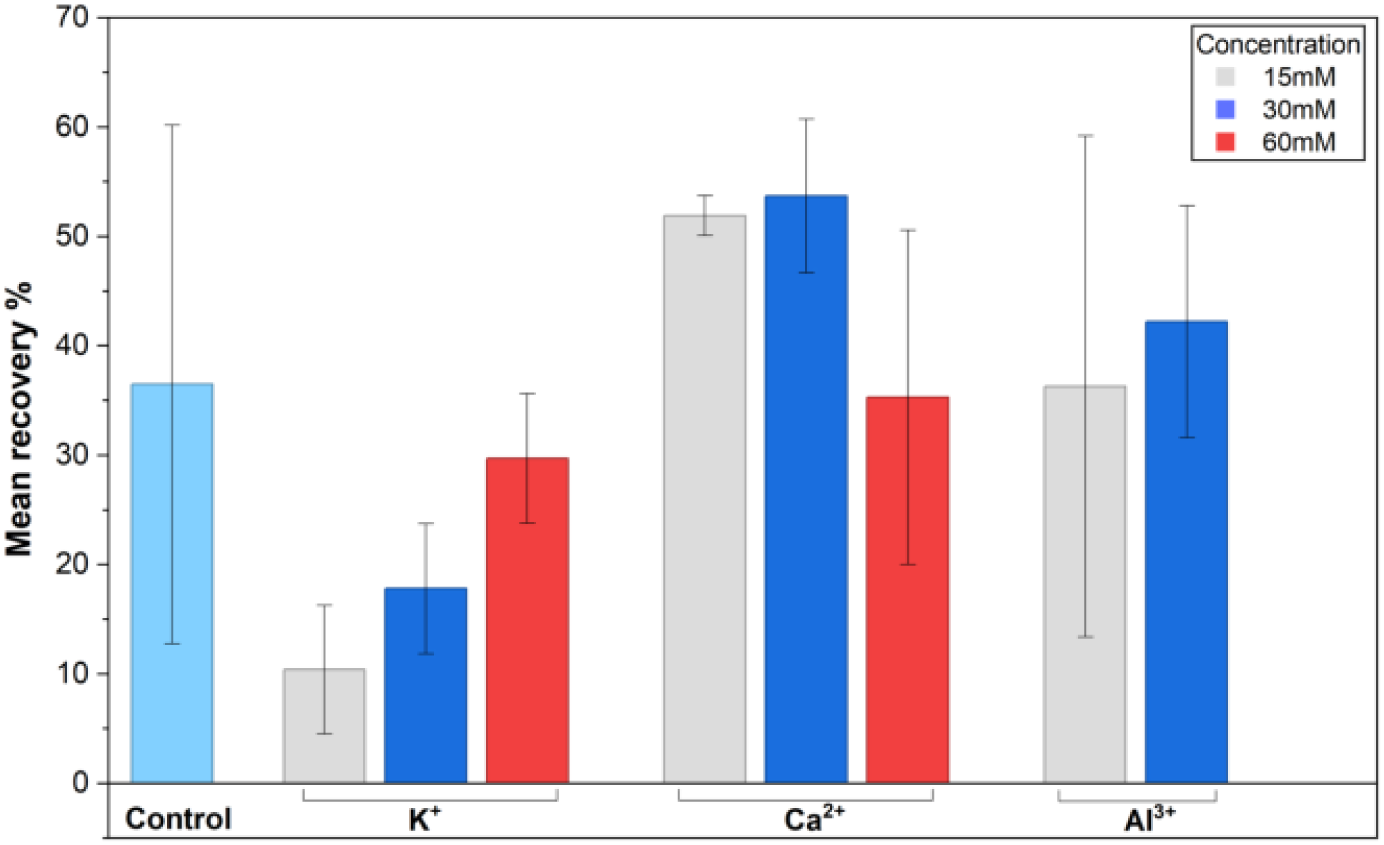
Mean recovery by ion charge and concentration

At 15 mM, Ca^2+^ systems exhibited the highest mean recovery (∼52%), significantly greater than both K^+^ (∼10%) and Al^3+^ (∼36%). This trend continued at 30 mM, with Ca^2+^ again maintaining the highest recovery (∼54%), followed by Al^3+^ (∼42%) and K^+^ (∼18%). At 60 mM, recovery decreased across all ion types, with Ca^2+^ dropping to ∼35%, and K^+^ to ∼30%. Post hoc comparisons confirmed that Ca^2+^ systems recovered significantly more than Al^3+^ (*p* = 0.011) and K^+^ (*p <* 0.001).

These results suggest that Ca^2+^ is most effective at supporting network recovery, consistent with its ability to stabilize *κ*CG junction zones while maintaining sufficient dynamics to permit reformation after deformation. The relatively low recovery for K^+^ systems indicates that while monovalent ions promote fast initial gelation, the resulting networks lack the robustness to recover efficiently, consistent with their role in helix stabilization without broader crosslinking. Al^3+^, despite inducing carboxylate shoulders in FTIR consistent with HA participation, showed variable recovery: moderate at 15–30 mM, but moduli could not be probed at 60 mM concentration due to network collapse. This collapse reflects over-screening since at 60mM, a much higher ionic strength is delivered indicates that the Al^3+^-mediated semi-IPN becomes brittle and over-crosslinked at high concentrations.

Thus, recovery behavior reinforces the FTIR and rheology results. K^+^ systems show no evidence of IPN formation and poor recovery, Ca^2+^ systems may form a semi-IPN due to the distinct binding mechanism but more clearly represent robust *κ*-CG dominated networks with high elastic moduli and recoverability, and Al^3+^ systems form a semi-IPN via HA co-crosslinking, but one that is prone to brittleness and failure at elevated ion concentrations.

These findings highlight the balance between ion valency, electrostatic screening, coordination chemistry, and recovery capacity in defining the functional character of HA-kCG gels.

## 4. Conclusion

This study demonstrates that interpenetrating network formation, structure and recovery of HA–*κ*CG networks are influenced by charge density and coordination behavior of the crosslinking ion. K^+^ stabilizes *κ*-CG helices without involving HA, producing relatively weak networks which exhibit poor recovery and no IPN formation. Ca^2+^ reinforces *κ*CG junction zones, yielding the highest elastic moduli and recovery capacity, with evidence that a semi-IPN may form. Al^3+^ introduces HA–carboxylate coordination in addition to sulfate interactions, confirming co-crosslinking and establishing a semi-IPN. However, this appears to result in brittle behavior and intermediate recovery profile. These results highlight the need to consider both ion charge and concentration in designing polysaccharide-based hydrogels, as they determine not only the strength of the network formed but also whether true interpenetration occurs and how efficiently the resulting network will recover under deformation. Although the recovery values obtained in this study can be further optimized to match more desirable recovery rates which are usually above 70%, we are able to understand from these findings that HA–*κ*CG networks enriched with Ca^2+^ seem to be the most promising candidate for injectable scaffolds or wound dressings that require robust recovery profile and with further optimization may be able to achieve self-healing properties

## Supporting Information

### 1. Additional Figures

**Figure S1:**
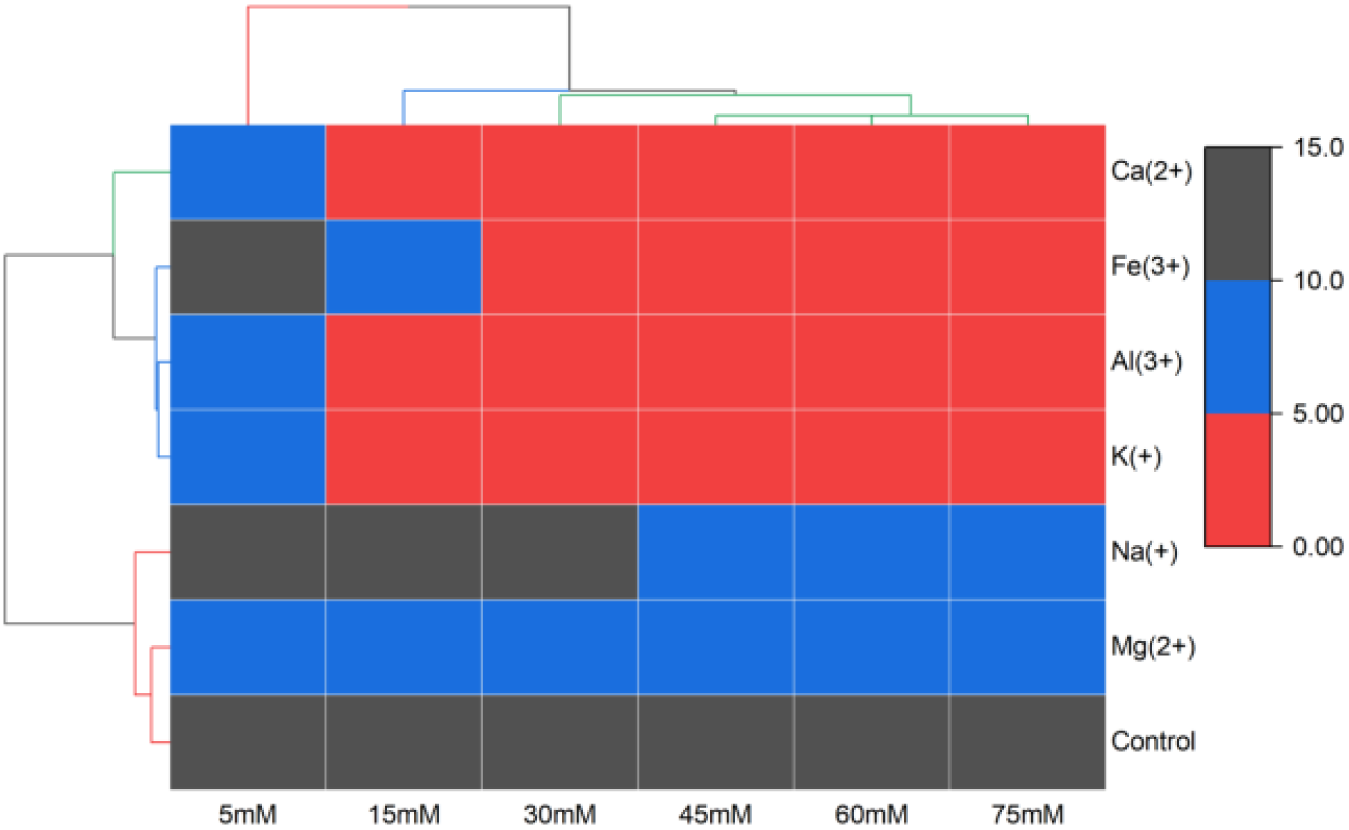
Heatmap with dendrogram showing hierarchical clustering of ion- and concentration-specific gelation trends

**Figure S2:**
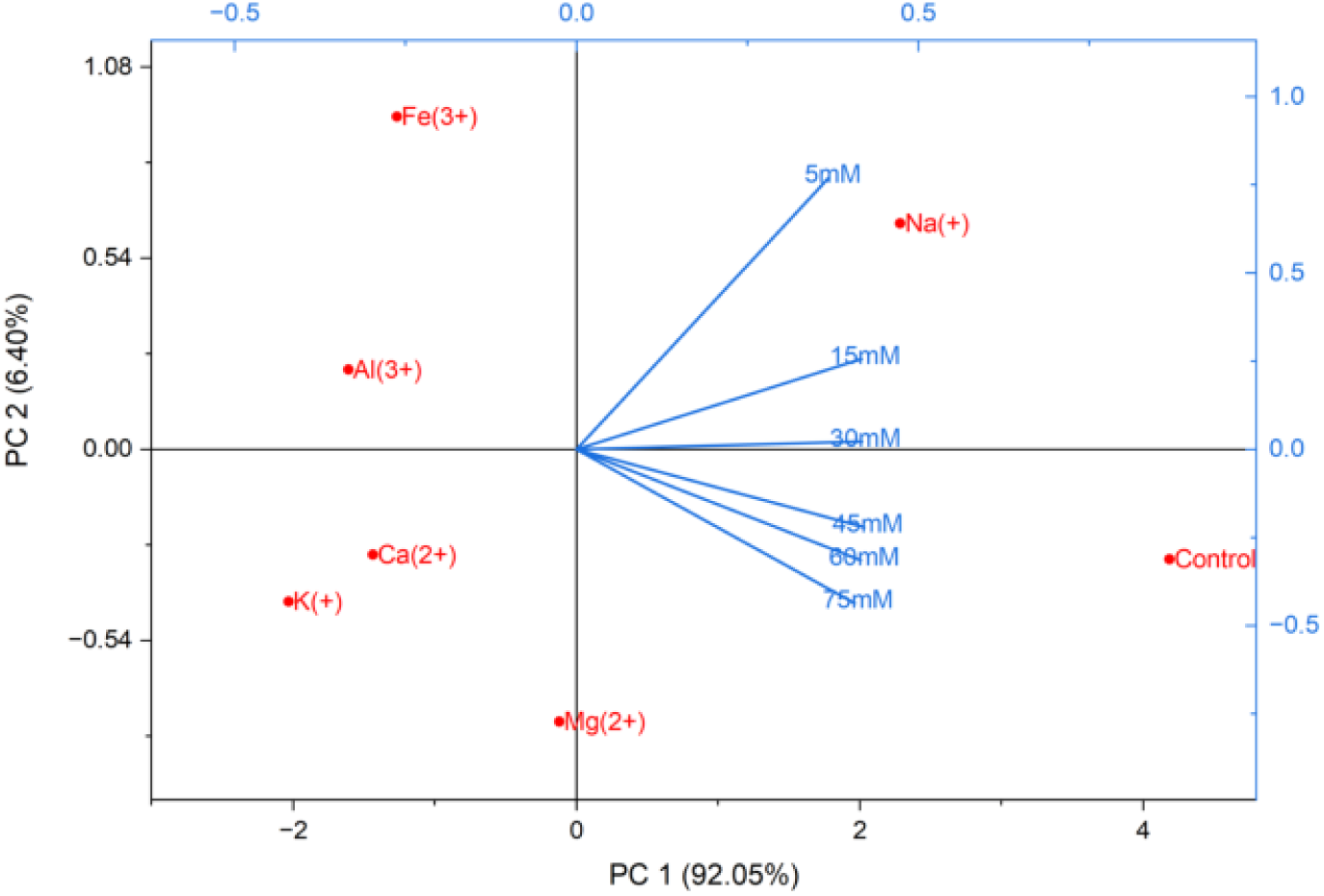
PCA scores plot showing principal component analysis of ion- and concentration-specific gelation trends

**Figure S3:**
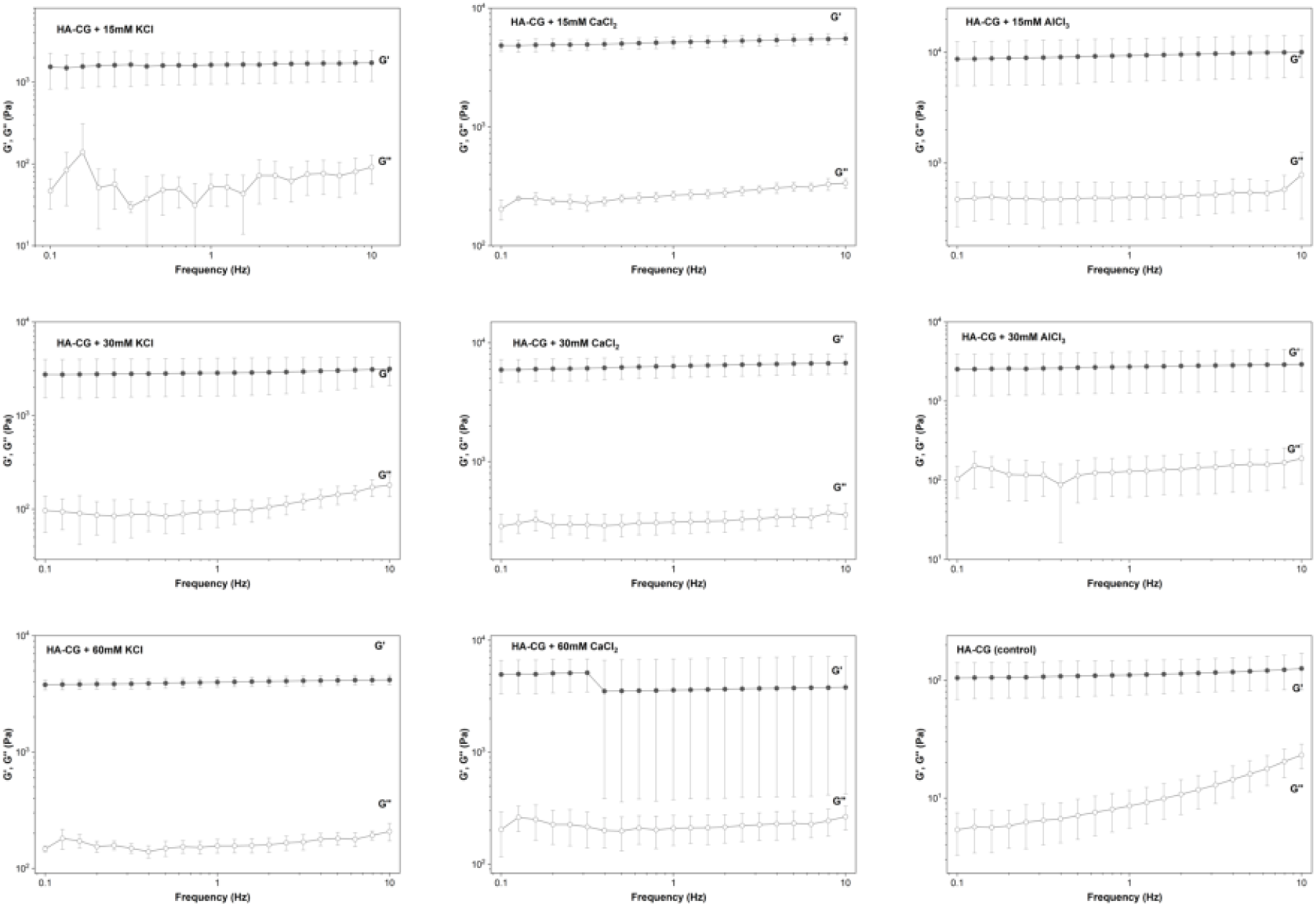
Frequency sweep plots across all ion conditions

**Figure S4:**
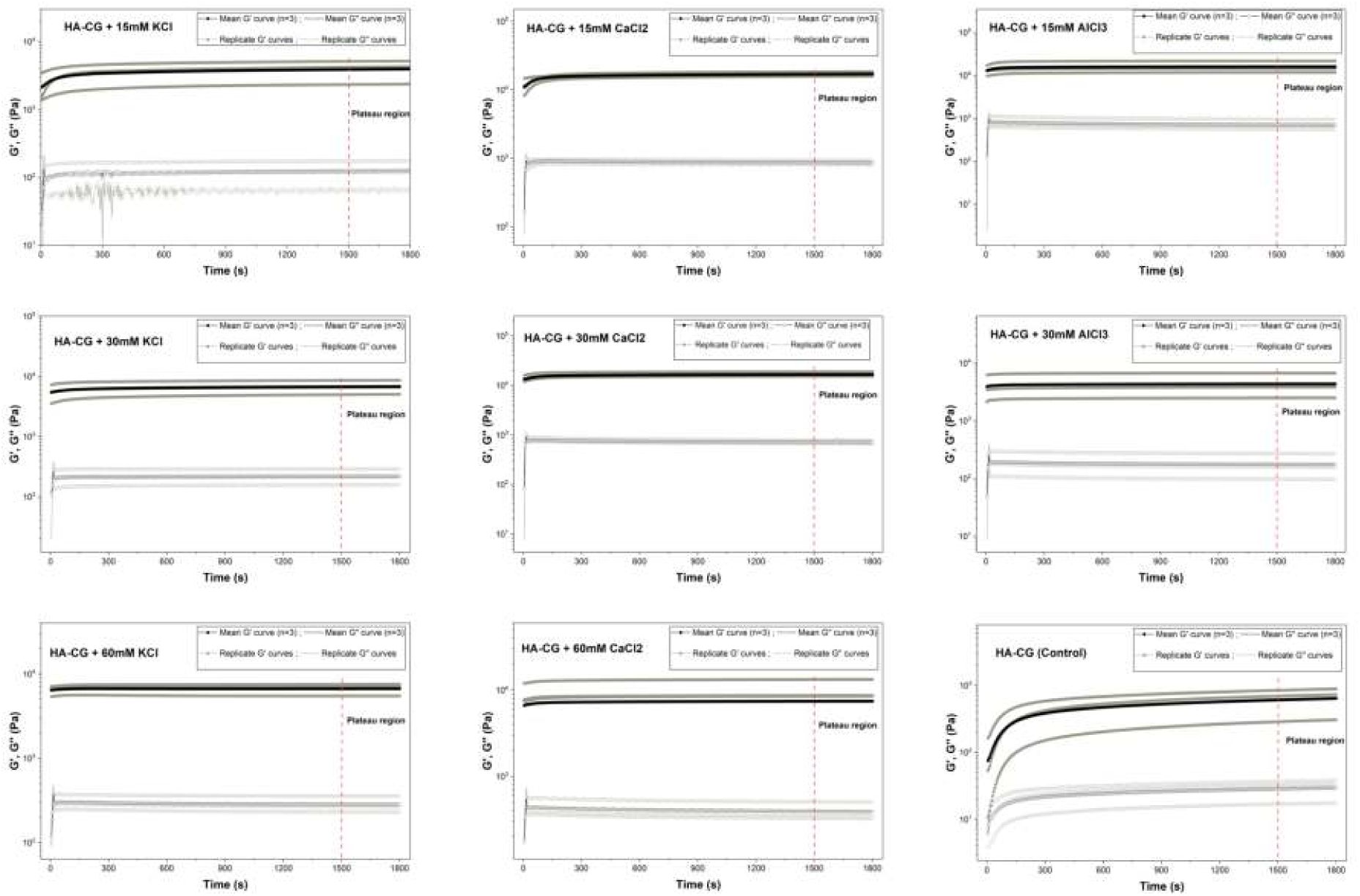
25^◦^C Timesweep plots across all ion conditions

**Figure S5:**
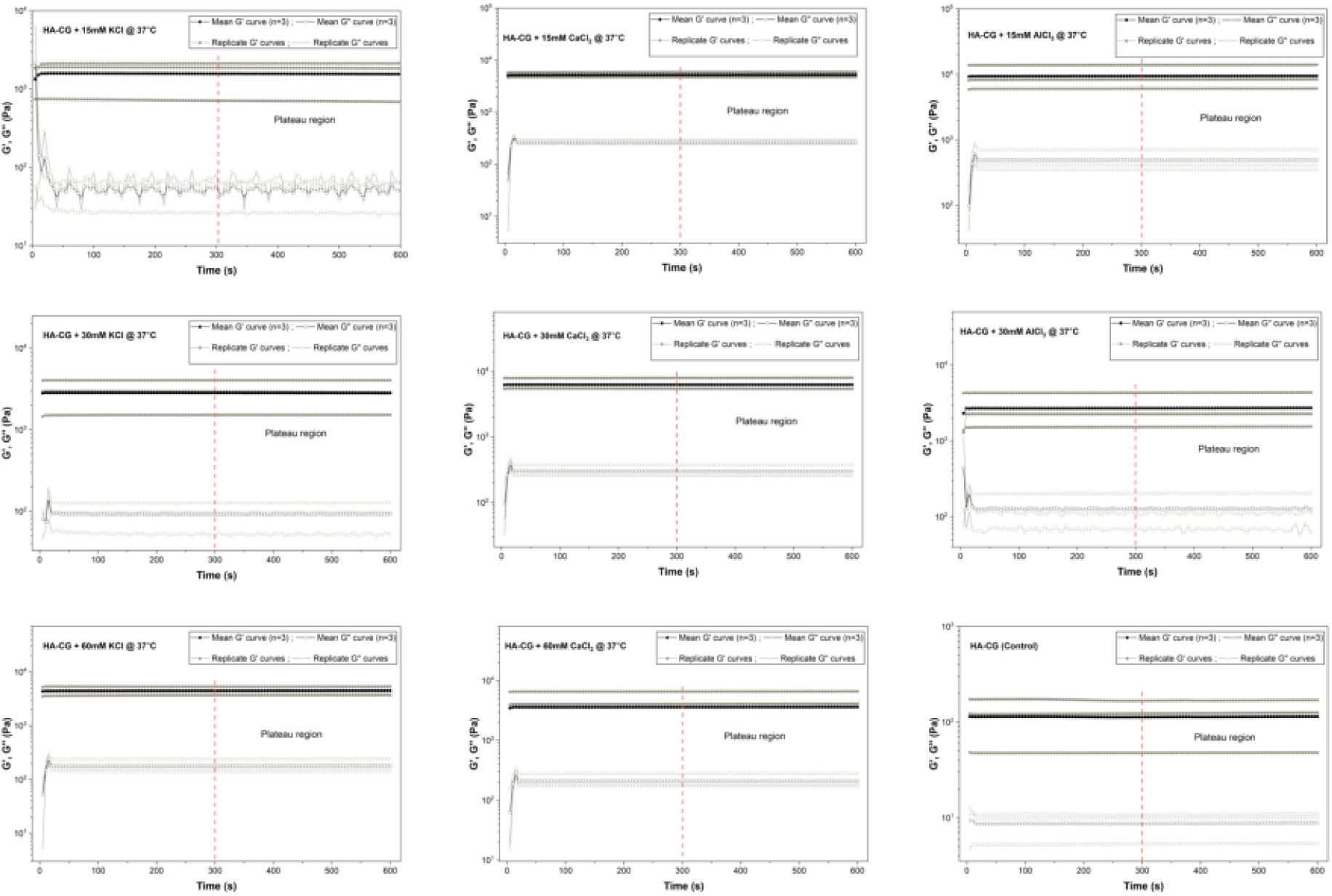
37^◦^C Timesweep plots across all ion conditions

**Figure S6:**
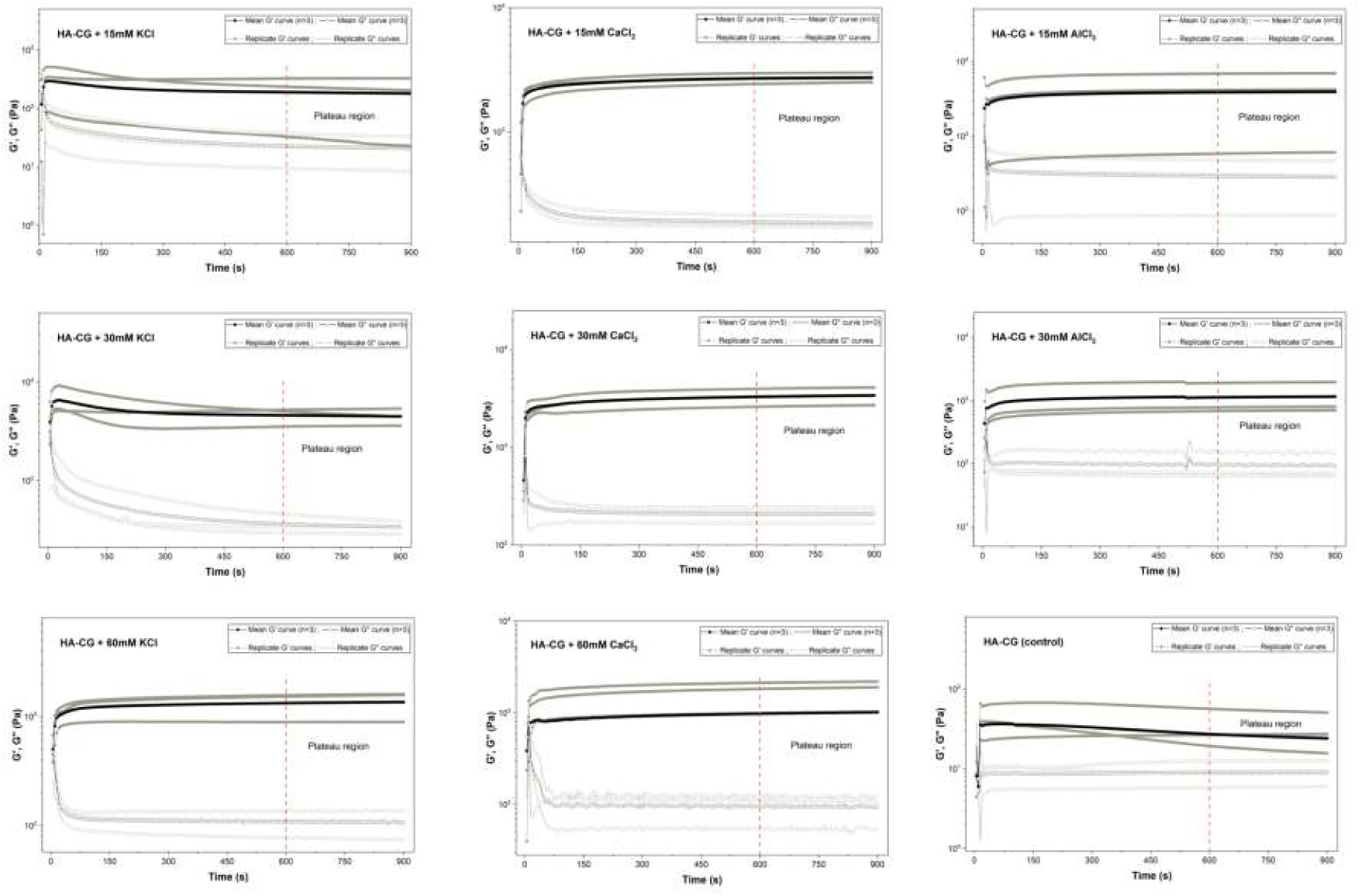
Recovery timesweep plots across all ion conditions

### Statistical Analysis Report (MANOVA)

The following results summarize descriptive statistics, assumption checks, omnibus tests, and post hoc analyses for the rheological outcomes considered in this study.

#### Descriptive Statistics (Mean ± SD)

**Table S1:** Descriptive statistics by ion valency and concentration.

| Ion (valency) | Conc. | $G'$ Loss % | $G'$ at 37 °C | Recovery (1 min) |
| --- | --- | --- | --- | --- |
| Al<sup>3+</sup> | 15 mM | 42.2 ± 6.5 | 9507.5 ± 4101.5 | 36.3 ± 22.9 |
| Al<sup>3+</sup> | 30 mM | 38.0 ± 2.8 | 2721.2 ± 1451.8 | 42.2 ± 10.6 |
| Al<sup>3+</sup> | 60 mM | 0.0 ± 0.0 | 0.0 ± 0.0 | 0.0 ± 0.0 |
| Ca<sup>2+</sup> | 15 mM | 69.0 ± 2.2 | 5255.8 ± 642.4 | 51.9 ± 1.8 |
| Ca<sup>2+</sup> | 30 mM | 62.2 ± 3.9 | 6302.7 ± 1463.0 | 53.7 ± 7.0 |
| Ca<sup>2+</sup> | 60 mM | 51.9 ± 1.3 | 4916.8 ± 1441.7 | 35.3 ± 15.3 |
| Control | 0 mM | 82.2 ± 1.7 | 113.9 ± 61.3 | 36.5 ± 23.7 |
| K<sup>+</sup> | 15 mM | 61.2 ± 10.4 | 1565.4 ± 760.1 | 10.4 ± 5.9 |
| K<sup>+</sup> | 30 mM | 82.2 ± 5.9 | 2830.2 ± 1268.0 | 17.8 ± 5.9 |
| K<sup>+</sup> | 60 mM | 32.8 ± 4.1 | 4499.6 ± 821.1 | 29.7 ± 5.9 |

#### Normality Tests

##### Univariate (Shapiro–Wilk)

*G*′_loss%_ : *W* = 0.933, *p* = 0.059 (approx. normal); *G*′_37°C_ : *W* = 0.902*, p* = 0.0096 (not normal); Recovery (1 min): *W* = 0.914*, p* = 0.0183 (not normal).

##### Multivariate (Mardia)

Skewness = 22.40*, p* = 0.013 (not normal); Kurtosis = 0.26*, p* = 0.796.

#### Multicollinearity Check (Pearson r)

No strong multicollinearity observed:

*G*′_loss%_ vs. *G*′_37°C_ : *r* = 0.009; *G*′_loss%_ vs. Recovery: *r* = 0.316; *G*′_37°C_ vs. Recovery: *r* = 0.498.

#### Homogeneity of Variances (Levene’s Test)

All *p >* 0.05; assumption met.

*G*′_loss%_ :*p* = 0.350; *G*′_37°C_ : *p* = 0.388; Recovery: *p* = 0.697.

#### MANOVA (Pillai’s Trace)

#### Univariate ANOVAs

All dependent variables (*G*′_loss%_, *G*′_37°C_, Recovery) show significant effects of IonValency, IonConc, and their interaction.

**Table S2:** MANOVA omnibus tests (Pillai).

| Effect | $F$ | $p$ |
| --- | --- | --- |
| IonValency | 11.09 | < 0.001 |
| IonConc | 9.08 | < 0.001 |
| Interaction | 7.44 | < 0.001 |

#### Effect Sizes (Partial η^2^)

*G*′_loss%_: 0.95 (Valency), 0.93 (Conc), 0.77 (Interaction);

*G*′_37°C_: 0.58, 0.32, 0.72;

Recovery (1 min): 0.55, 0.28, 0.49.

#### Tukey Post Hoc (Recovery by IonValency)

Ca^2+^ *>* Al^3+^: *p* = 0.011; K^+^ *<* Ca^2+^: *p <* 0.001.

#### MANOVA

##### MANOVA (Pillai)

IonValency: Pillai = 1.87, *F* = 11.09, *p* = 0; Ion-Conc: Pillai = 1.18, *F* = 9.08, *p* = 0; Interaction: Pillai = 1.79, *F* = 7.44, *p* = 0.

